# The two-component system YesMN promotes pneumococcal host-to-host transmission, and regulates genes involved in zinc homeostasis

**DOI:** 10.1101/2022.08.29.505782

**Authors:** M. Ammar Zafar, Alicia Costa-Terryl, Taylor M. Young

## Abstract

The ability to sense and respond rapidly to the dynamic environment of the upper respiratory tract (URT) makes *Streptococcus pneumoniae* (*Spn*) a highly successful human pathogen. Two-component systems (TCS) of *Spn* sense and respond to multiple signals it encounters allowing *Spn* to adapt and thrive in various host sites. *Spn* TCS have been implicated in their ability to promote pneumococcal colonization of the URT and virulence. As the disease state can be a dead-end for a pathogen, we considered whether TCS would contribute to pneumococcal transmission. Herein, we determined the role of YesMN, an understudied TCS of *Spn*, and observe that YesMN contributes towards pneumococcal shedding and transmission but is not essential for colonization. The YesMN regulon includes genes involved in zinc homeostasis and glycan metabolism, which are upregulated during reduced zinc availability in a YesMN dependent fashion. Thus, we identify the YesMN regulon and the molecular signals it senses that lead to the activation of genes involved in zinc homeostasis and glycan metabolism. Furthermore, in contract to *Spn* mono-infection, we demonstrate that YesMN is critical for high pneumococcal density in the URT during influenza A (IAV) coinfection. We attribute reduced colonization of the *yesMN* mutant due to increased association with and clearance by the mucus covering the URT epithelial surface. Thus, our results highlight the dynamic interactions that occur between *Spn* and IAV in the URT, and the role that TCS play in modulation of these interactions.

## Introduction

The nasopharynx is the initial site of acquisition and the primary reservoir of the human-adapted extracellular respiratory pathogen *Streptococcus pneumoniae* (*Spn*; the pneumococcus). *Spn* resides in the mucosal surface of the upper respiratory tract (URT), and carriage is typically asymptomatic. From this niche in the URT, pneumococcus can spread to another host or cause disease within the same host. Disease states associated with *Spn* include pneumonia, septicemia, otitis media, and meningitis. Invasive pneumococcal disease is associated with over a million deaths worldwide (1-3).

The ability to colonize and thrive in different host environments is a hallmark of *Spn* pathogenesis. Multiple studies have shown that pneumococcus modifies its global gene expression depending on the colonization site (4-7). Adapting and responding to the dynamic environments that bacterial pathogens encounter requires rapid responses, some of which are mediated through two- component signal transduction systems (TCSs). Bacterial TCS generally consists of two proteins, a histidine kinase (HK) that senses external stimuli, and a response regulator (RR). Upon autophosphorylation of the HK in response to external stimuli, the phosphate group is transferred to the RR, which undergoes a conformation change allowing it to act as a regulator (8, 9).

The *Spn* genome encodes 13 TCS, each consisting of an HK and a cognate RR, and one orphan response regulator. As with other bacterial pathogens, several of these TCS contribute to pneumococcal colonization and virulence. They sense a variety of signals, including, but not limited to, reactive oxygen species, oxidative stress, antibiotics, antimicrobial peptides, pH, and temperature. In response, they modulate the expression of genes involved in reducing oxidative stress, uptake of nutrients, carbon metabolism, modulating competence and fratricide, releasing bacteriocins, and other factors that affect host-pathogen interactions. Several TCS of *Spn* have been extensively characterized, including BlpRH, ComDE, and CiaRH. BlpRH is involved in quorum sensing and bacteriocin production; ComDE, through quorum sensing, regulates *Spn* transformation machinery under specific environmental conditions; CiaRH regulates expression of the serine protease HtrA, the stress response machinery, and negatively influences competence (8, 10). Furthermore, there is crosstalk signaling between pneumococcal TCS, where they respond to the same environmental signal (ComDE and CiaRH both respond to acid stress) and regulate the expression of each other (10-12).

Specific pneumococcal factors, such as pneumolysin (13), *dlt* locus dependent d-alanylation of *Spn* teichoic acid (14), and the type and amount of *Spn* capsular polysaccharide (15), are essential for *Spn* host-to-host transmission. These factors modulate the host inflammatory response or the dynamic interactions with the mucus protecting the URT epithelial layer. Recent high-throughput transposon mutagenesis (Tn-seq) screens have identified *Spn* factors important for colonization, virulence, and host-to-host transmission (14, 16, 17). As TCS can regulate hundreds of genes, it was not surprising that they would be identified in these screens to contribute toward pneumococcal pathogenesis. However, one of the least characterized TCS, YesMN (also called TCS07), was found to contribute toward pneumococcal host-to-host transmission in two dissimilar Tn-seq approaches with different animal models (14, 17), indicating that YesMN regulates factors that are important determinants of pneumococcal spread.

The putative regulon of YesMN was initially identified by overexpressing YesN (RR) and performing next-generation RNA sequencing (18). Genes involved in host-derived glycan metabolism and transport were among the YesMN regulon. Specifically, genes encoding enzymes for N-glycan metabolism and transport were identified to be upregulated in a YesMN dependent manner (18). These results were unsurprising, as free carbon is scarce in the URT (19). *Spn* devotes considerable resources to acquiring carbon by cleaving host-glycans, such as heavily glycosylated mucin proteins that form the mucus layer (20, 21). Nevertheless, the signal and the biological relevance in the nasopharynx of YesMN remain poorly defined.

Herein we aimed to validate the *in vivo* role of YesMN suggested by Tn-seq screens with a *yesMN* deletion mutant tested outside of the context of a mixed infection. In addition, we sought to better define the YesMN regulon using a *yesMN* deletion strain rather than over-expression. Our results indicate that YesMN does indeed play a role in transmission but not in URT colonization. Transcriptomic analysis revealed the YesMN regulon includes zinc homeostasis genes and validated the published role in glycan metabolism (18). Importantly, we found that YesMN impacts its regulon in response to changes in zinc levels, suggesting that zinc may be the signal that is sensed by YesMN.

## Results

### Effect of *yesMN* deletion on pneumococcal colonization, shedding and host-to-host transmission

A transposon mutagenesis screen revealed putative *Spn* factors that contribute toward pneumococcal nasopharyngeal colonization and shedding (14, 16). A candidate of particular interest was the HK YesM that forms part of the two-component system (TCS) YesMN (TCS07) (**Fig 1A**). Overexpression of the response regulator (RR) YesN in D39 (Type 2 isolate) leads to an increase in the expression of genes that code for N-linked glycan-metabolizing enzymes. Also, compared to the parental D39, *ΔyesMN* had a slightly reduced virulence in a lung infection model (18).

**Figure 1.**
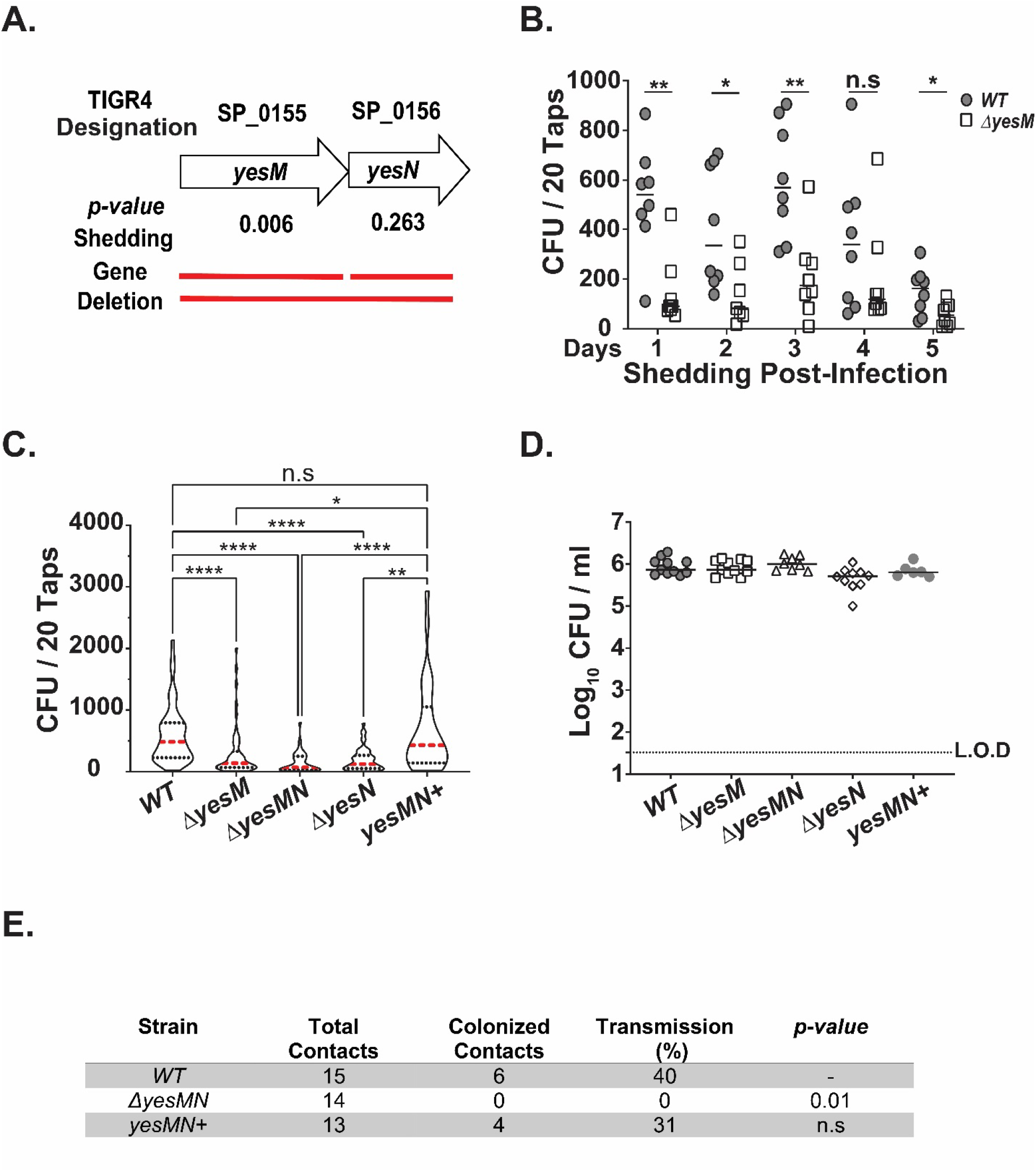
The two-component system YesMN of pneumococcus contributes towards its shedding and host-to-host transmission. **A**. Schematic representation of the genetic organization of the TCS YesMN of *S. pneumoniae* with the TIGR4 designation. Adjusted *P* values for shedding for each gene are listed, with the solid line representing the gene deletions constructed for the current study. **B**. 4 day old pups were challenged i.n. with the indicated strain, and nasal secretions were collected and bacterial load was enumerated for 5 days post-infection. Indicated are median CFU values for each day, and each symbol represents the number of CFU measured from a single pup (n≥5 for each strain tested). Shedding differences between the two strains on a given day were calculated using the Mann-Whitney *U* test. **C**. Pups were challenged i.n. at day 4 of age with the indicated construct, and bacterial shedding from nasal secretions quantified for 5 days post-infection. Indicated are violin plots of pooled 5 days of shedding with median shown in red. Data are from 6 to 10 pups per group. Statistical differences were calculated using the Kruskal-Wallis analysis of variance with Dunn’s posttest. **D**. Colonization density of each pneumococcal construct in cultures of URT lavage fluids obtained from pups at day 9 of age. Median values are shown. Dotted line represents the limit of detection (L.O.D). **E**. Summary of transmission efficiency of the WT, *ΔyesMN* mutant and the corrected strain (*yesMN*^*+*^) from colonized index pups at the age of 4 days to naïve contact pups in the same litter. The transmission efficiency was determined by the number of contact pups (at a 1:1 ration to index mice) colonized by S. pneumoniae at the age of 14 days. Fisher’s exact test was used to determine the statistical difference between the WT and the *ΔyesMN* mutant and *yesMN*^*+*^. *, *P ≤* 0.05; **, *P* ≤ 0.01; ****, *P ≤* 0.0001, n.s, not significant.

We were particularly intrigued by identifying the *yesMN* TCS in our shedding screen as of the 13 pneumococcal TCS, YesMN regulation is poorly understood (8). Moreover, another *in vivo* transposon screen to identify putative transmission factors of *Spn* using a ferret model identified the RR YesN as being necessary for host-to-host transmission (17). To confirm that the TCS YesMN impacts pneumococcal shedding, we constructed mutants with unmarked, in-frame deletion of HK *yesM* (*ΔyesM*), the RR *yesN* (*ΔyesN*), or both *yesMN* (*ΔyesMN*). Deletion of *yesMN* was not associated with impaired growth (18) and all the constructed mutants in the study grew as WT type 4 isolate (data not shown).

To determine if YesMN contributes towards *Spn* shedding, we first assessed *ΔyesM*, which was identified in our shedding screen (14). Shedding was measured daily (Days 1 to 5 post-inoculation) from the inoculated pups. Compared to the wild-type (WT) type 4 strain, the isogenic mutant *ΔyesM* shed poorly over five days (**Fig. 1B**). Because of established day-to-day variation (22), daily shedding values were also pooled for comparison (**Fig 1C**). The HK (*ΔyesM*), RR (*ΔyesN*) mutants, and the complete TCS knockout (*ΔyesMN*) all showed reduced median shedding compared to either the WT type 4 strain or the corrected mutant (*yesMN*+ mutant). These results included an intra-litter comparison of strains inoculated into pups from multiple litters to control for host and environmental effects on shedding. The contribution of *yesMN* to shedding was not due to differences in colonization density, as all the strains colonized equally well at the conclusion of the shedding experiment at the age of nine days (**Fig. 1D**).

Next, we determined whether reduced shedding attributed to the *yesMN* mutants manifests as a defect in host-to-host transmission. Using a 1:1 ratio of *Spn* inoculated pups (index) to uninoculated littermates (contact pups), ∼40 percent transmission was observed with the WT type 4 strain. In contrast, no transmission events were detected with the *ΔyesMN* strain. Transmission to WT levels was restored with the corrected strain (*yesMN*^+^) (**Fig. 1E**). Furthermore, all constructs colonized the pups at a high density at the conclusion of the transmission study at the age of 14 days (**Fig. S1**). Thus, we validated our and others’ observations, confirming that under *Spn* mono-infection, YesMN does not affect colonization but is required for sufficient pneumococcal shedding to allow for host-to-host transmission.

### Comparative transcription analysis of YESMN mutant

Recently, RR *yesN* overexpression led to the identification of the YesMN regulon (18). As pneumococcal TCS exist in a balanced crosstalk, overexpression of a single RR can potentially affect other signaling pathways. To provide more physiologically relevant molecular testing conditions, we determined the global gene expression patterns in a *ΔyesMN* strain to identify the putative YesMN regulon. The WT type 4 and the *ΔyesMN* strains were grown in TSY medium, and total RNA transcript abundance was measured using RNA-seq. A total of 134 differentially regulated genes were identified in the *ΔyesMN* compared to the WT strain (Log2FC>±1, Padj<0.05), of which 88 were down-regulated and 49 were upregulated (**Fig. 2A-C**) (**Supplemental Table 1**). Besides genes of unknown function, most of the genes identified were involved in the uptake of nutrients and their metabolism (**Fig. 2B-C**). The top downregulated genes encode proteins involved in zinc homeostasis, host-glycan and carbohydrate transport and metabolism, and the locus associated with purine metabolism (**Table 1**). The putative YesMN regulon included genes previously shown to be necessary for pneumococcal colonization and or shedding (14, 16). Notably, genes that encode enzymes involved in host glycan metabolism (**Table 1**) identified in our RNA-seq analysis were correspondingly identified when YesN was ectopically overexpressed in the D39 background (18). Thus, our RNA-seq results align with the previously described study and identify new members of the YesMN regulon. Moreover, examination of the upstream region of the genes that form the YesMN regulon did not yield a consensus binding site for the rr07. Furthermore, CiaR that forms the part of the CiaRH TCS and several of its regulon members were in the list of genes whose transcripts were upregulated (∼2 fold) in the *ΔYesMN* strain (**Supplemental Table S1**).

**Table 1.**
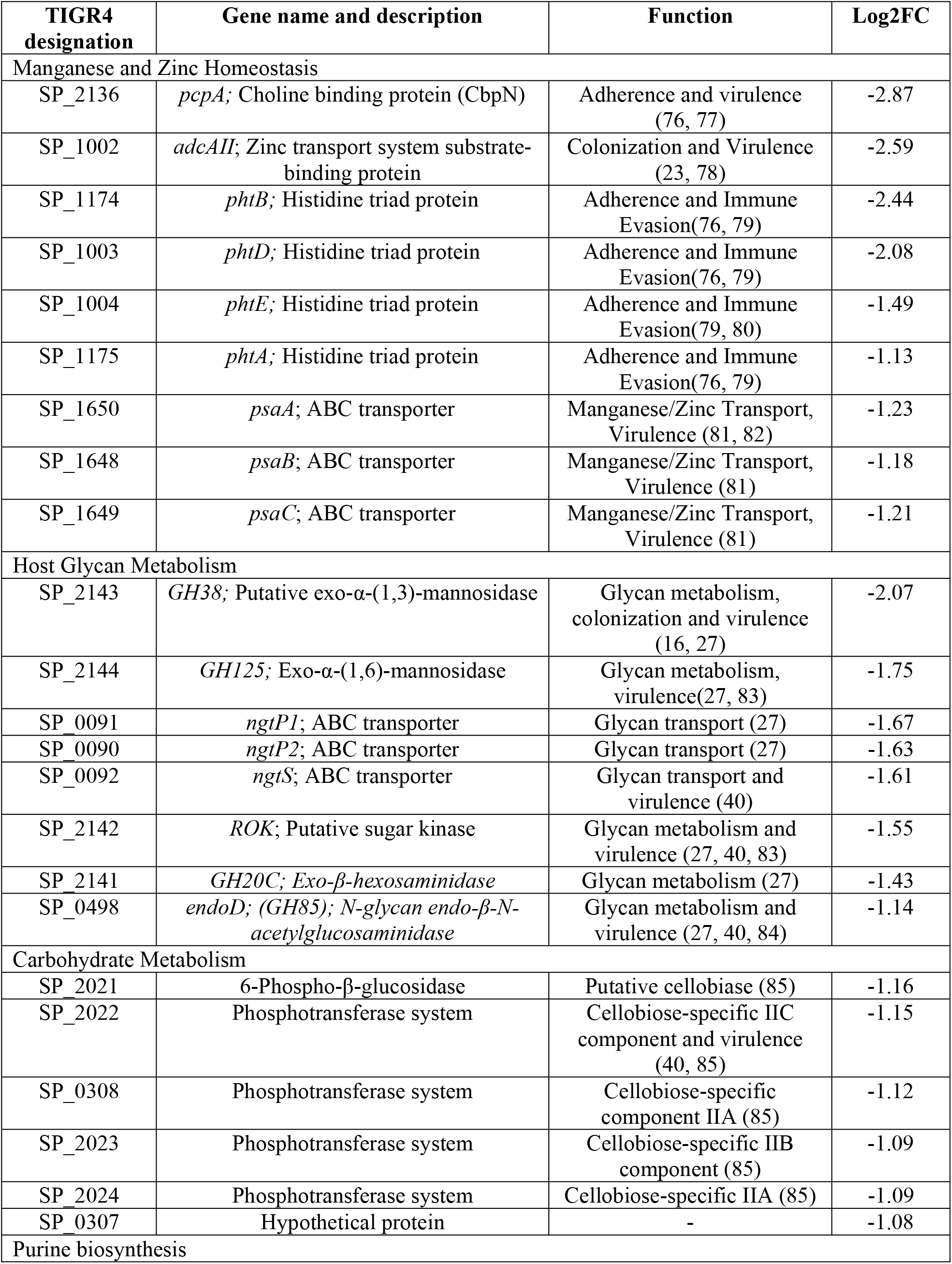
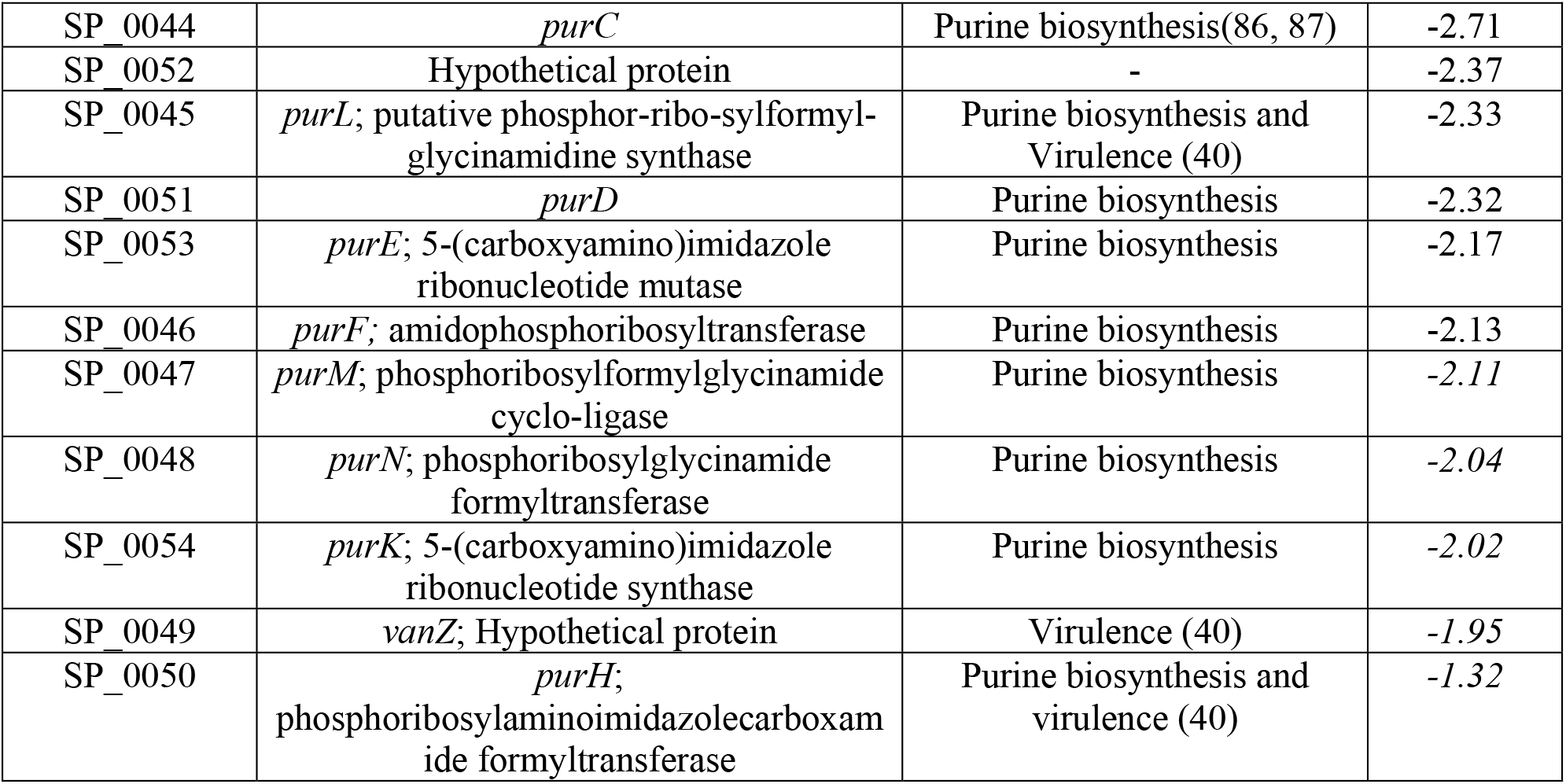
Major functional classes of genes whose transcripts were downregulated in a *ΔyesMN* mutant. Gene name, description and function were based on information retrieved from KEGG and uniprot databases, and literature searches.

**Figure 2.**
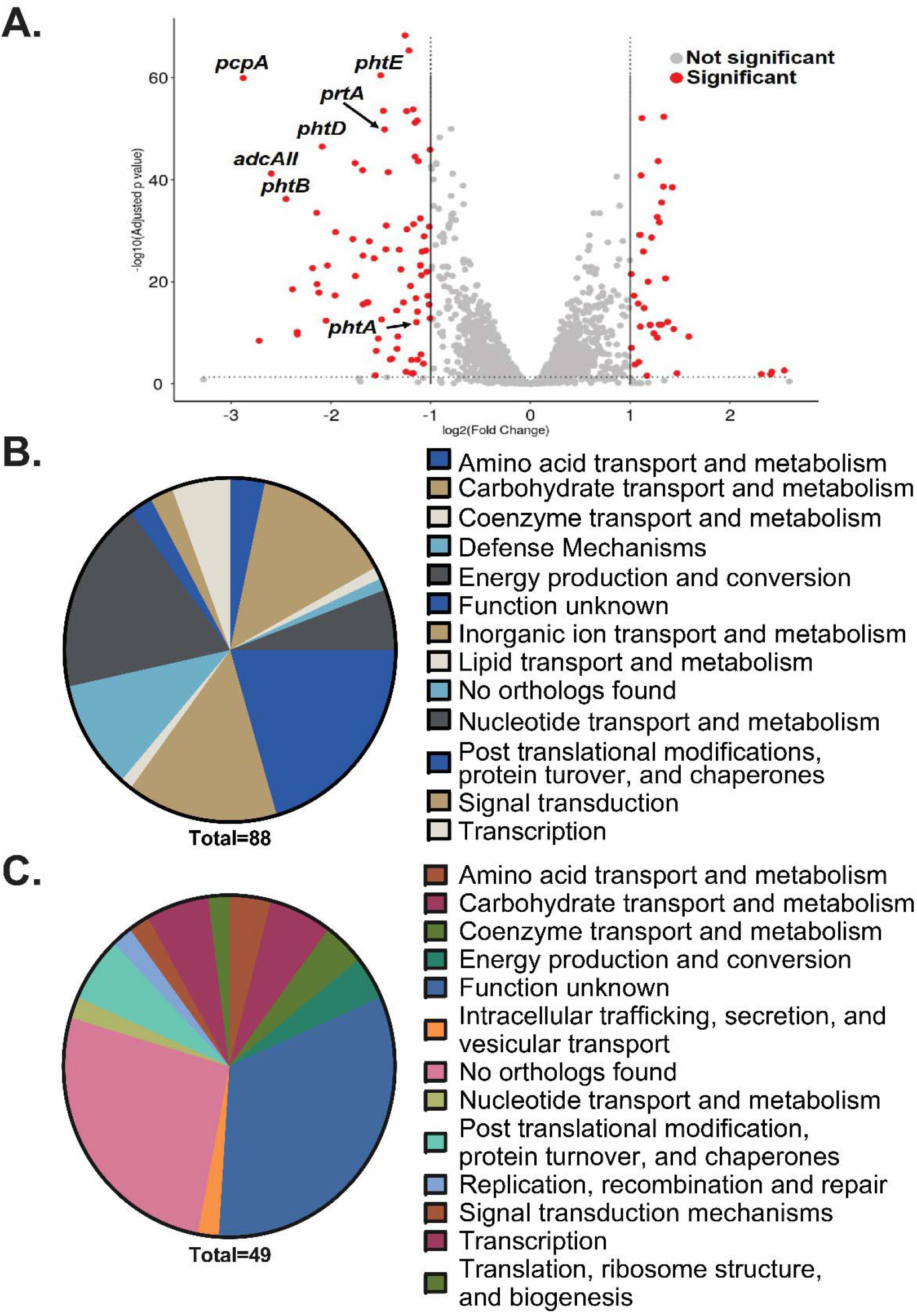
Identification of the genes regulated by the pneumococcal TCS YesMN. The transcriptome of wild-type TIGR4 and the isogenic *ΔyesMN* mutant was compared through RNA-seq analysis. Strains were grown in TSY to OD_620_= 0.4, RNA isolated and prepared for analysis as described in Materials and Methods section. **A**. Volcano plot depicting genes whose RNA transcripts were differentially regulated (Log2FC> ±1; *P* < 0.05). Genes involved in Zinc homeostasis are highlighted. **B-C**. Functional annotation of genes whose transcripts in the *ΔyesMN* mutant compared to the wild-type were either **B**. down-regulated **C**. or upregulated.

To validate the results from the RNA-seq analysis, quantitative real-time PCR was performed targeting 11 specific genes implicated in either zinc homeostasis or host-glycan metabolism (**Table 1**). Neuraminidase A (*nanA*) and B (*nanB*) were selected as control as they were not identified by the RNA-seq screen to be modulated by YesMN. The YesMN regulated genes to be validated included *adcAII*, which encodes an adhesion lipoprotein with specificity for Zn^2+^ (23); *phtD* and *phtE* known as pneumococcal histidine triad (Pht) proteins that form an operon with *adcaII*. Pht family proteins are also involved in Zn^2+^ acquisition, and upregulated under low Zn^2+^ (24, 25). *pcpA* that encodes for choline-binding protein N was also observed in the RNA-seq screen to be regulated by YesMN. However, in the presence of Mn^2+^, PsaR represses the expression of *pcpA* (26). Besides, genes involved in Zn^2+^ homeostasis, we tested *endoD*, GH20C, ROK, GH38, and GH125, which are involved in N-glycan and fetuin metabolism (18, 27).We observed a statistically significant ∼2-fold reduction in the transcripts of the selected genes when comparing *ΔyesMN* to the WT strain (**Fig. 3A-B**). No reduction was observed in the transcripts of control genes (*nanA* and *nanB*) that do not belong to the YesMN regulon (**Fig. 3A**). Taken together, our results strongly suggest that YesMN regulates genes involved in zinc homeostasis and glycan metabolism.

**Figure 3.**
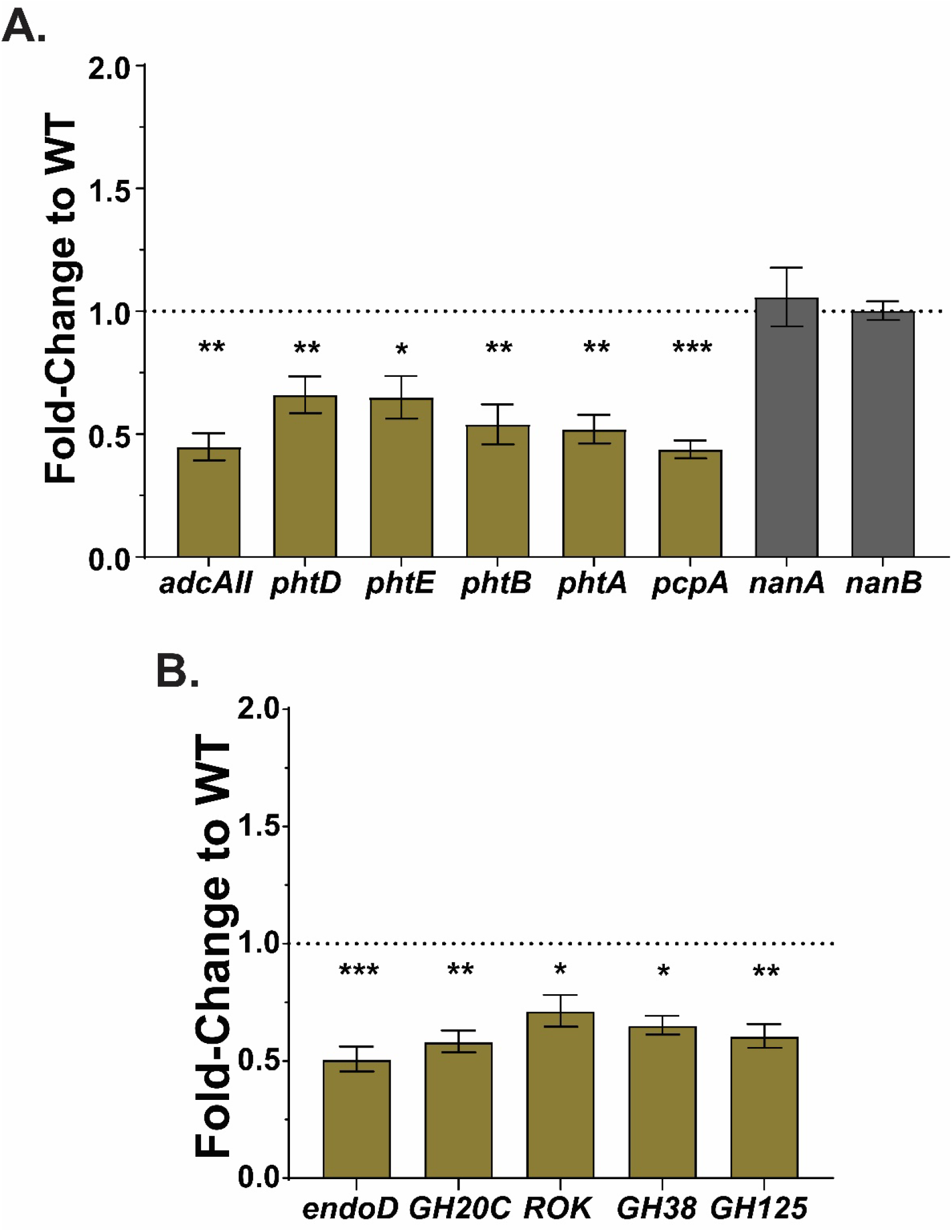
YesMN regulates genes involved in zinc homeostasis and glycan metabolism. Differential gene expression in *ΔyesMN* of selected genes by quantitative real-time PCR (qRT-PCR). WT and the *ΔyesMN* isogenic mutant were grown to OD_620_= 0.4 in TSY broth at 37°C, RNA isolated and prepared for qRT-PCR analysis. Shown is fold-change in transcription of genes between WT and *ΔyesMN* involved either in **A**. zinc homeostasis or **B**. glycan metabolism. *nanA* and *nanB* that are not part of the YesMN regulon were tested as controls (grey bars). TIGR4 *16s* (*rrsA, SP_0775*) specific primers were used as the housekeeping gene for 2^*-ΔΔC*^ _*T*_ analysis. For each biological replicate (n ≥ 3 for WT and *ΔyesMN*), qRT- PCR was conducted in triplicate. Bars represent mean ± SEM. Statistical differences between the WT and the *yesMN* mutant were calculated using the Mann-Whitney *U* test. *, *P* ≤ 0.05; **, *P* ≤ 0.01; ***, *P*≤ 0.001.

### YesMN appears to sense Zn^2+^ to modulate its regulon

Based on our RNA-seq analysis that identified genes involved in Zn^2+^ homeostasis belonging to the YesMN regulon, we hypothesized that YesMN responds to Zn^2+^ level in the environment. Previously, *adcaII* and *pht* genes transcripts were found to be upregulated when grown in N, N, N’, N’- tetrakis-(2-pyridylmethyl)-ethylene-diamine (TPEN), a membrane-permeable zinc chelator (28). Correspondingly, TSY growing WT strain when shifted to TSY containing 30 µM TPEN led to an upregulation in the transcript levels of the *adcaII, pht* genes, as well as *pcpA* (∼15 to 30 fold) (**Fig. 4A**). In addition, transcripts of glycan metabolism genes were upregulated (**Fig. 4B**), albeit less (∼2 to 4 fold) than the zinc homeostasis genes. In contrast, in the *ΔyesMN* strain, TPEN treatment did not increase the transcripts of genes involved in zinc homeostasis or glycan metabolism to the same level as WT (**Fig. 4A-B**). Zinc homeostasis genes were still upregulated (∼3-4 fold) in the *ΔyesMN* strain under TPEN, suggesting that other factors also contribute to the increase in expression of these genes under reduced levels of Zn^2+^. Collectively, our data suggest that the Zn^2+^ homeostasis and glycan metabolism genes form part of the YesMN regulon and respond to reduced Zn^2+^ levels.

**Figure 4.**
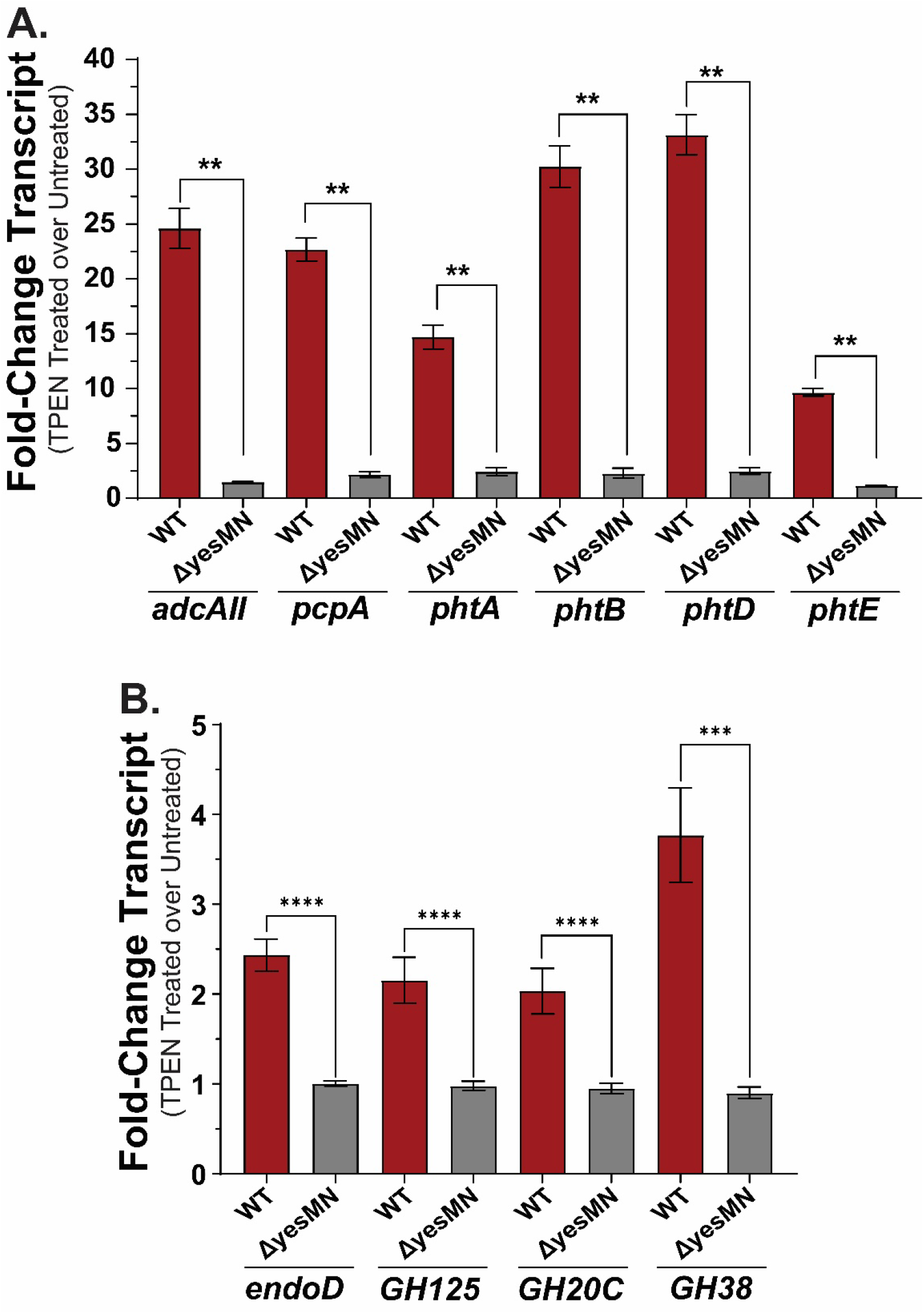
YesMN responds to reduced availability of zinc. Quantitative real-time PCR analysis was used to examine gene expression in the WT and the *ΔyesMN* isogenic mutant in response to zinc chelation. Strains were grown to OD_620_= 0.4 in TSY broth at 37°C. Half of the culture was transferred to a clean tube and treated with TPEN and the other half remained untreated; these were grown for additional 30 min at 37°C. RNA was isolated and prepared for qRT-PCR analysis as described in M&M. TIGR4 *16s* (*rrsA, SP_0775*) specific primers were used as the housekeeping gene for 2^*-ΔΔC*^_*T*_ analysis. Bars (mean ± SEM) in red represent WT whereas grey bars represent *ΔyesMN*, and indicate the fold-change in transcript abundance between treated (TPEN) or untreated of **A**. zinc homeostasis genes or **B**. glycan metabolism genes. For each biological replicate (n ≥ 3 for WT and *ΔyesMN*), qRT-PCR was conducted in triplicate. Statistical differences between the WT and the *ΔyesMN* mutant were calculated using the Mann-Whitney *U* test. **, *P* ≤ 0.01; ***, *P*≤ 0.001; ****, *P ≤* 0.0001.

### YesMN is required for robust colonization during concurrent influenza A infection

Epidemiological data show that bacterial-influenza <u>A v</u>irus (IAV) infections occur in more than 30% of all cases, predominantly with pneumococcus (29, 30). These coinfections are associated with increased morbidity and mortality (31, 32). *Spn*-IAV coinfection alters the nasopharyngeal landscape of the infected host, with an increase in inflammation and mucus production, allowing *Spn* to proliferate further (33-35). Recently, a transposon mutagenesis screen using a type 3 isolate identified *Spn* factors important for nasopharyngeal colonization with a concurrent IAV infection. The list of genes identified included several that form part of the YesMN regulon and are involved in N-glycan metabolism. (36). We hypothesized that even though *ΔyesMN* strain colonizes as WT under mono-infection, its regulon members are potentially critical for colonization during IAV coinfection.

To test our hypothesis, we inoculated pups with either the *ΔyesMN* strain or the WT type 4 isolate at four days of life and, on day 9, gave them the mouse adapted Influenza A virus/Puerto Rico/8/34(H1N1), hereafter referred to as “PR8” and followed the modulation of *Spn* shedding from the pups over five days post-IAV infection. Analogous to our previous studies, the WT strain shed at a high level during IAV coinfection (22). Similar to the results for mono-infection (**Fig. 1C**), the *ΔyesMN* strain also had a defect in shedding during IAV coinfection (**Fig. 5A**). However, unlike mono-infection, where the *ΔyesMN* strain colonized as WT (**Fig. S1**), with a concurrent IAV infection the mutant colonized poorly (**Fig. 5B**), suggesting that the YesMN regulon is critical for robust colonization in the changing landscape of the nasopharynx. Furthermore, the shedding and colonization defect was YesMN specific as the *yesMN*^+^ strain shed and colonized as WT. Thus, our data shows that even though YesMN is dispensable during mono-infection, it is critical for pneumococcal colonization during an IAV coinfection and provides further evidence that *Spn*-IAV interactions are not necessarily synergistic in the upper respiratory tract.

**Figure 5.**
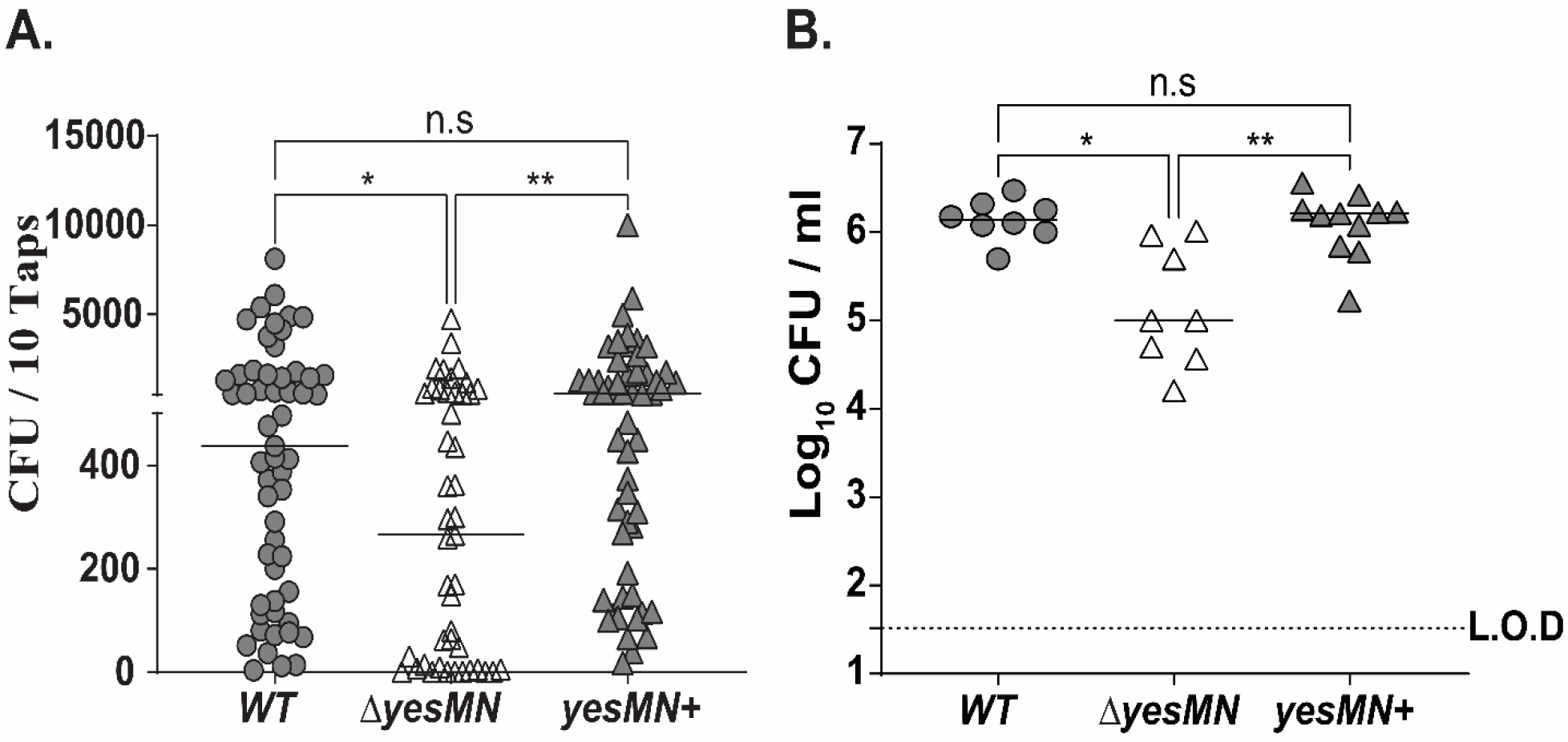
YesMN is required for robust pneumococcal shedding and colonization in the setting of influenza A coinfection. **A**. Four day old pups were challenged i.n. with either the WT, the isogenic yesMN mutant, or the corrected *yesMN*^*+*^ strain. Five days later, pups were challenged with influenza A virus, and pneumococcal shedding in nasal secretions was quantified daily for five days. Shown is pooled median shedding values of 5 days, where each symbol represents CFU values obtained from a single pup. **B**. Colonization density of each pneumococcal construct in cultures of URT lavage fluids obtained from pups at day 14 of age. Median values are shown. Dotted line represents the limit of detection (L.O.D). Data are from 8 to 11 pups per group. Statistical differences were calculated using the Kruskal-Wallis analysis of variance with Dunn’s posttest. *, *P ≤* 0.05; **, *P* ≤ 0.01; n.s, not significant.

### YesMN impacts the interaction of pneumococcus with human nasal fluid

The YesMN regulon includes genes that encode proteins that act as adhesins, promote virulence and contribute toward N-linked glycan metabolism (**Table 1**). Previous studies demonstrated that high Pneumococcal shedding is dependent upon (**i**.) ability of *Spn* to cause a robust inflammatory response in the URT that results in an increase in nasal secretions and (**ii**.) its ability to escape mucus present on the URT epithelial surface (13, 15, 33, 37, 38). We therefore examined whether the reduced shedding from the URT of the *ΔyesMN* strain during mono-infection is due to low level inflammation. The chemokines (CXCL-1, CXCL-2) and cytokine interleukin-1β (IL-1β) are considered hallmarks for *Spn*-induced inflammation in the URT (13), and their levels were measured following inoculation with WT or the *yesMN* mutant. At 2 days post-inoculation, we did not observe upregulation of these inflammatory markers in mice inoculated with either strain. (**Fig. S2**). Thus our qRT-PCR data likely precludes *Spn* mediated inflammatory response in the URT as the basis for reduced shedding associated with the *ΔyesMN* mutant.

Based on our observation, we postulated that an increase in nasal secretions leads to a reduced load of the *ΔyesMN* mutant in URT during coinfection. To determine whether YesMN contributes to interactions with mucus, we performed a previously described *in vitro* solid-phase human mucus binding assay using pooled <u>h</u>uman <u>n</u>asal fluid (hNF) (39). Adherence of the *ΔyesMN* strain to mucus was significantly higher compared to the WT type 4 isolate, suggesting that YesMN encodes proteins that promote mucus evasion (**Fig. 6**). This enhanced adherence was not because of reduced capsular polysaccharide (CPS), as both the mutant and the WT strain made similar levels of CPS (**Fig. S3**). Moreover, this enhanced interaction with hNF was specific to *ΔyesMN* mutant as the *yesMN*^+^ strain restored reduced interaction with hNF. Overall, our results demonstrate that YesMN regulated factors that influence interactions with the protein that constitute hNF.

**Figure 6.**
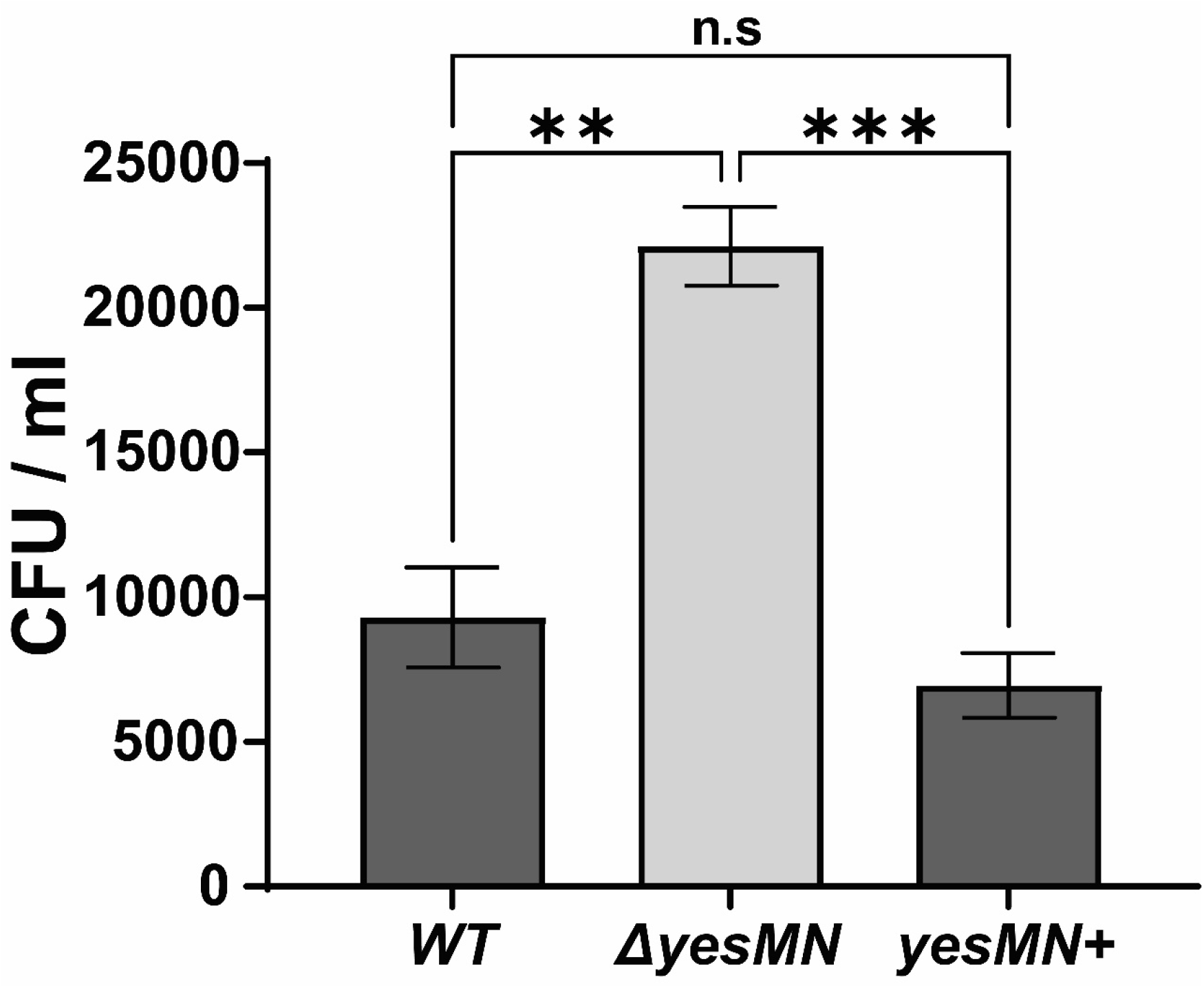
YesMN affects human nasal fluid association. Comparison of level binding of T4S, the *ΔyesMN* mutant and the *yesMN*^+^ strain to immobilized human nasal fluid (hNF). Bacteria (2x 10^4^ CFU/100 μl DMEM) were incubated with 100 μl of 10 µg of immobilized hNF in presence of 0.1% BSA for 2hr at 30°C. After 19 washes, adherent bacteria were determined by resuspending with 0.001% Triton X-100 following plating on TSY agar plates supplemented with 200 μg/ml streptomycin. Values are means of three independent determinations in triplicate ± SEM. Statistical differences were calculated using the Kruskal-Wallis analysis of variance with Dunn’s posttest. **, *P* < 0.01; ***, *P* < 0.001; n.s, not significant.

## Discussion

In the current study, we sought to increase our understanding of nasopharyngeal colonization and host-to-host transmission of the human-adapted pathogen *Streptococcus pneumoniae* using an infant mouse model (22). Two-component systems respond to various overlapping signals that allow for quick adaptation and establishment by *Spn* within the nutrient-poor but dynamic environment of the nasopharynx (8). Two independent Tn-seq screens identified one of the least characterized TCS (YesMN) as contributing to the spread of *Spn* from one host to another without impacting its colonization of the URT (14, 17). Thus, to provide a molecular understanding of how YesMN contributes towards pneumococcal spread, we identified its regulon members, and the signal it uses to modulate their expression.

Previous studies with adult mice demonstrated that YesMN does not necessarily impact nasopharyngeal colonization but did have reduced virulence (18, 40, 41). Our tractable infant mouse model allowed us to examine the contribution of YesMN in pneumococcal colonization, shedding, and host-to-host transmission. YesMN knockout strains had reduced shedding without impacting colonization (**Fig. 1**). Consequently, we determined whether YesMN impacts *Spn* host-to-host transmission as our previous work had shown that reduced shedding correlates with poor transmission dynamics (13). Our transmission studies demonstrated that a *ΔyesMN* strain is unable to transmit, providing further evidence that reduced shedding impacts transmissibility, and highlights the importance of YesMN to pneumococcal spread to another host. Hence, our data established that YesMN-coordinated regulation of the genes in the nasopharynx is critical for robust *Spn* shedding and transmission To gain insight into the genes within the YesMN regulon that could influence the virulence and transmission phenotypes, we performed RNA-seq with a serotype 4 WT (TIGR4) and isogenic Δ *yesMN* strains. We observed that transcripts of genes involved in Zn^2+^ homeostasis, N-linked glycan metabolism, carbohydrate metabolism, and purine metabolism were downregulated in the *ΔyesMN* strain (**Table 1**).

This data aligns well with the transcript changes published by Andreassen *et al*. using the genetically distinct D39 strain (serotype 2) (18). Although *Spn* isolates are genetically diverse with over 90 serotypes, the genes within the YesMN regulon are highly conserved across *Spn* isolates (18, 42). An important caveat to this study is that the RNA-seq was performed on bacteria grown in TSY, which is likely to be nutritionally quite different than the nasopharyngeal environment. However, several of the genes identified in this *in vitro* screen have been previously identified as essential for either colonization or shedding (14, 16, 40), suggesting that the TSY growth medium is not inappropriate for such analyses Cross-talk between different TCSs can affect the identification of environmental signals that they respond to activate their specific regulon. In our RNA-seq analysis, we identified genes that are involved in zinc homeostasis, which we verified through subsequent qRT-PCR and illustrated that YesMN contributes under Zn^2+^ chelation conditions to the upregulation of zinc homeostasis and glycan metabolism genes. Trace metals, including zinc, are critical for the enzymatic function of both eukaryotic and prokaryotic cells (43). Zinc deficiency, modeled in murine and human studies, is associated with the reduced phagocytic activity of polymorphonuclear cells (PMN) and degradation of the mucus layer, which leads to increased disease severity and bacterial burden (44-49). Likewise, an increase in dietary zinc is concomitant with a reduction in invasive pneumococcal disease (50-52).

In keeping with the critical role of zinc, the host modulates Zn^2+^ levels in the blood and respiratory tract in response to pneumococcal infection (Nutritional Immunity) (47, 53). In this dynamic micro-niche environment where Zn^2+^ concentrations can fluctuate, *Spn* has to strike a delicate balance between Zn^2+^ sufficiency and toxicity, maintained by Zn^2+^ homeostasis genes (53-56). The URT is a nutritionally poor environment, with low levels of Zn^2+^ where the host mucin proteins, which contain labile Zn^2+^, potentially serve as a source (57, 58). Correspondingly, *Ogunniyi et al*. (59) observed an upregulation in the URT of the transcripts of the histidine triad proteins (*pht*). These proteins, together with AdcA and AdcaII, are required for zinc uptake (23, 25).

The HK YesM is predicted to contain a large extracellular loop allowing it to detect small molecules and ions, and providing it with the ability to respond to a diverse number of signals (8). Interestingly, when we used TPEN, a high-affinity chelator of Zn^2+^, to mimic the zinc starvation that *Spn* likely encounters in the URT, we observed a YesMN-dependent regulation of the *pht, adcaII*, and glycan metabolism genes (**Fig. 4A-B**). Thus, our data suggest that YesMN coordinates the expression of these genes, resulting in enhanced shedding and subsequent transmission of *Spn*. Surprisingly, we identified *pcpA* to be a member of the YesMN regulon. In contrast to the *adcaII* and the *pht* genes, whose transcripts are upregulated under reduced Zn^2+^ levels in the environment, *pcpA* expression is regulated by CodY (60) and PsaR (26). PsaR regulatory roles are dependent on divalent ion concentration. High levels of Mn^2+^ cause downregulation, whereas high levels of Zn^2+^ lead to upregulation of *pcpA* (61). Generally, there are higher levels of Mn^2+^ in the nasopharynx compared to Zn^2+^ (26, 61); therefore, PsaR would likely downregulate the expression of *pcpA*. However, *pcpA* expression in the URT is higher compared to the blood (4), and functional antibodies have been observed in children without local or systemic infection (62), suggesting that *pcpA* is expressed in the URT. The opposing role of PsaR and YesMN would suggest the essentiality of *pcpA* expression under specific conditions, and that YesMN under reduced levels of Zn^2+^ might override the downregulation of *pcpA* by PsaR.

As *Spn*-IAV coinfections occur frequently and can result in higher morbidity and mortality (31, 32), we investigated the role of YesMN during *Spn*-IAV coinfection of the nasopharynx. Similar to *Spn* mono-infection, we observed reduced *Spn* shedding from pups colonized with the *ΔyesMN* mutant (**Fig. 5A**). However, in contrast to mono-infection, pups colonized with the *ΔyesMN* mutant coinfected with IAV had a defect in URT colonization (**Fig. 5B**). During *Spn*-IAV coinfection, *Spn* shedding and URT colonization density is known to increase over time, which has been attributed to an increase in nasal secretions and reduced mucociliary action (33, 34, 63). Moreover, increased *Spn* shedding leads to an increase in host-to-host transmission (34). These data suggest that *Spn*-IAV coinfection is beneficial for *Spn* dissemination. However, *Spn* impacts IAV titers and host-to-host transmission negatively (64, 65). Furthermore, IAV infection leads to the dispersal of *Spn* biofilms and modulation of its transcriptome (66). Collectively, these data suggest a complex and possibly antagonistic relationship. Our results further build on these data, where the YesMN regulon provides *Spn* with an advantage in the dynamic environment of the URT during *Spn*-IAV coinfection. Additionally, our hNF binding assay suggests that YesMN regulated factors contribute towards the ability to escape nasal secretion and not be cleared by the host. Thus, it is likely that YesMN dependent changes in gene expression allow *Spn* to navigate the inflamed URT, where N-linked glycan metabolism and other factors allow *Spn* to thrive and escape in high numbers to transmit to another host.

In conclusion, our studies have enhanced our understanding of one of the least characterized TCS of *S. pneumoniae*. We demonstrate that YesMN contributes towards *Spn* shedding and host-to-host transmission during pneumococcal mono-infection. We attribute reduced shedding of the *ΔyesMN* mutant to an inability to escape the mucus that lines the epithelial layer of the URT, suggesting that YesMN encodes factors critical for *Spn*-mucus interactions. These YesMN-dependent factors include surface proteins required for zinc homeostasis and glycan metabolism. Furthermore, we provide evidence for antagonistic interaction during *Spn*-IAV coinfection, where YesMN promotes the ability of *Spn* to thrive in the inflamed URT co-colonized with IAV. Elucidation of specific YesMN dependent factors that impact the ability of *Spn* to persist and thrive in the inflamed URT would provide further mechanistic insights into bacterial colonization and dissemination.

## Methods

### Ethics Statement

This study was conducted according to the guidelines outlined by National Science Foundation animal welfare requirements and the *Public Health Service Policy on Humane Care and Use of Laboratory Animals* (67). All animal work was done according to the guidelines provided by the American Association for Laboratory Animal Science (AALAS) (68) and with the approval of the Wake Forest Baptist Medical Center Institutional Animal Care and Use Committee (IACUC). The approved protocol number for this project is A20-057.

### Growth Conditions and Strain Construction

Strains used in the current study are listed in **Table 2**. Pneumococcal strains were grown at 37°C in tryptic soy broth (Becton Dickinson [BD]) supplemented with 0.3% yeast extract (TSY). The strains in this study are derivatives of AZ57, which is a streptomycin-resistant derivative of the type 4 strain TIGR4 (22). Once *Spn* cultures reached the desired optical density at 620 nm (OD_620_), they were washed, and diluted in sterile phosphate-buffer saline (PBS) for intranasal inoculation and other assays. To quantitate *S. pneumoniae*, serial dilutions were routinely plated on TSY agar-streptomycin (200 µg/ml), supplemented with either catalase (6,300 U/plate; Worthington Biochemical Corporation) or 5% defibrinated sheep’s blood, and incubated overnight at 37°C with 5% CO_2_. The *yesM, yesMN* and *yesN* deletion mutants were constructed using the Janus cassette (69) in a two-step process. Briefly, the Janus cassette was amplified from genomic DNA isolated from strain AZ59 (13) using the MasterPure DNA purification kit (Lucigen), with flanking regions (∼1kb) upstream and downstream of the gene of interest using isothermal assembly. Strain AZ57 was then transformed with the PCR product and the transformants selected on TSY agar with kanamycin (125 µg/ml). The resulting mutant was confirmed through PCR, and selected for making an in-frame clean gene deletion. The mutant was transformed with PCR product using primers listed in **Table S2**. All in-frame clean gene deletions contain the first and last 5 amino-acid coding sequence of the gene of interest. The corrected mutant *yesMN*^*+*^ was constructed using primers (YesMN-US-1/YesMN-DS-6) on genomic DNA from AZ57 and the resultant PCR product was transformed into strain AZ83, and transformants selected on TSY agar-streptomycin (200 µg/ml).

**Table 2.** Strains used in the study.

| Strain | Description<sup>a</sup> | Reference or Source |
| --- | --- | --- |
| AZ57 | P2406; (TIGR4Sm<sup>R</sup>) | (22) |
| AZ59 | P2423; TIGR4 *ply::kan-rpsL*<sup>+</sup>; Km<sup>R</sup>,Sm<sup>S</sup> | (13) |
| AZ69 | AZ57; *yesM::kan-rpsL*<sup>+</sup>; Km<sup>R</sup>,Sm<sup>S</sup> | This Study |
| AZ81 | AZ69; Δ*yesM*; Sm<sup>R</sup> | This Study |
| AZ83 | AZ57; *yesMN::kan-rpsL*<sup>+</sup>; Km<sup>R</sup>,Sm<sup>S</sup> | This Study |
| AZ84 | AZ83; Δ*yesMN*; Sm<sup>R</sup> | This Study |
| AZ85 | AZ57; *yesN::kan-rpsL*<sup>+</sup>; Km<sup>R</sup>,Sm<sup>S</sup> | This Study |
| AZ86 | AZ86; Δ*yesN*; Sm<sup>S</sup> | This Study |
| AZ88 | AZ83; *yesMN*<sup>+</sup>, Sm<sup>R</sup> | This Study |

<sup>a</sup> Km, kanamycin; Sm, streptomycin.

### Infant mouse infection model for shedding, colonization and transmission

All mouse studies were performed with C57BL/6J Specific Pathogen Free (SPF) mice that were initially obtained from the Jackson Laboratory (Bar Harbor, ME), then bred and maintained in the animal facility at Biotech Place, Wake Forest Baptist Medical Center. The pups were housed with their dam (mother) for the duration of the study, and gained weight like uninfected animals.

Pups were infected as described previously (22). Briefly, at day 4 of life, the pups were inoculated intranasally (i.n.) with ∼2,000 CFU of *S. pneumoniae* suspended in 3 µl of PBS. Post-inoculation, shedding was determined by gently tapping the nares of pups (20 taps/pup) on to a TSY agar plate supplemented with either streptomycin (200 µg/ml) or kanamycin (125 µg/ml) to only allow for the growth of *Spn*. The nasal secretions were spread using a sterile cotton-tipped swab, and the plates incubated as described above. Because of inherent variability in shedding from day-to-day, the CFU values obtained from individual pups were pooled together for the 5 day duration of the study. To control for environmental and microbiome variability, shedding of different strains was compared for pups within the same litter. The mouse adapted Influenza A virus PR8 was used for co-infection studies. 5 days post inoculation with *Spn*, pups were challenged i.n with 2 × 10^4^ PFU of PR8 suspended in 3 µl of PBS. Daily shedding was quantified as above, but because of higher shedding values, only 10 taps/pup were carried out.

To determine colonization density, pups were euthanized at the designated time points by CO_2_ asphyxiation followed by cardiac puncture. The upper respiratory tract (URT) was exposed and lavaged with 200 µl of sterile PBS from a needle inserted into the trachea, and fluid collected from the nares. The limit of detection for *Spn* in lavage fluid was 33 CFU/ml.

Pneumococcal monoinfection transmission studies were performed as previously described (22). Briefly, at day 4 of life, half of a cohort of pups were randomly selected to be inoculated with the indicated *Spn* strain. These index mice were then returned to their uninfected littermates (contact pups) and cohoused for 10 days post-inoculation. To determine transmission from the index pups, all pups were euthanized at the age of 14 days, and nasal lavage fluid was collected and serially diluted on TSY-agar plates supplemented with appropriate antibiotic.

### RNA-seq Sample Preparation and Analysis

*Spn* cultures were grown as described above to an OD_620_ ∼0.4, and RNA isolated using Qiagen All prep bacterial DNA/RNA/protein kit following the manufacturer’s protocol. Purified RNA samples were analyzed for quality by electrophoretic tracing (Agilent Bioanalyzer). The Illumina TruSeq Stranded total RNA LT kit with Bacterial Ribo Depletion Probes (Illumina) was used to construct cDNA libraries from samples with RNA Integrity (RIN) values equal to 10. Briefly, 500 ng of total RNA was rRNA depleted, fragmented, reverse-transcribed to cDNA, then purified via AMPure XP magnetic beads. Sequencing adaptors were then ligated to end-repaired fragments, and the libraries were pre-amplified by PCR and quantified on a Qubit 3.0 (Thermo Fisher, USA). Dual indexed libraries were pooled and sequenced to a target read depth of 50M reads on an Illumina NextSeq 500 using 75 bp single-end sequencing and a high-output flow cell. For all samples, >90% of sequences achieved >Q30 Phred quality scores (FASTQC analysis, Babraham Bioinformatics). Contaminating adapters were removed with Trimmomatic (70).

Reads were aligned to the genomic reference *Streptococcus pneumoniae* TIGR4 using Bowtie 2 (71). Gene counts were determined using featureCounts software (72). Differentially expressed genes were identified by negative binomial modeling using DESeq2 (73) with false discovery correction (Benjamini-Hochberg). This work was performed by the Cancer Genomics Shared Resource of the Wake Forest Baptist Comprehensive Cancer Center.

### RNA isolation and qRT-PCR

Host RNA was isolated by lavaging the URT of pups inoculated with *Spn* or mock inoculated (PBS) at the designated time-point using RLT lysis buffer (Qiagen) to obtain RNA from the epithelium. RNA was purified and cDNA generated as per manufacturer’s (Bio-rad) instructions. Quantitative reverse- transcription-PCR (qRT-PCR) was performed using Bio-rad SYBR green master mix, ∼2.5ng cDNA, and 0.5 mM primers per reaction mix. Primers directed towards GAPDH (glyceraldehyde-3-phosphate dehydrogenase) were used as an internal control. Samples were run on CFX384 Touch real-time PCR detection system (Bio-Rad) in duplicate, and each run was repeated twice. RNA expression was quantified using the *ΔΔC*_*T*_ threshold cycle (*C*_*T*_) method. Primers used in the current study were previously described (74, 75).

qRT-PCR was performed on TSY-broth grown *Spn* cultures to confirm the genes that form the YesMN regulon. The WT and *ΔyesMN* strains were grown in TSY-broth statically at 37°C until they reached an OD_620_ ∼0.4 and [2 ml] was spun down at room-temperature at 15,500 x *g* for 2 min. Samples were treated with RNAprotect Bacteria Reagent (Qiagen), and RNA was isolated using the All prep bacterial DNA/RNA/protein kit following the manufacturer’s (Qiagen) protocol. cDNA and qRT-PCR were carried out as described above with the primer sets listed in **Table S3**.

To determine the effect of Zn^2+^ on the expression of YesMN-dependent genes, WT and *ΔyesMN* were grown statically in TSY broth at 37°C till they reached an OD_620_=∼0.3. At that point samples were divided and either treated with Zn^2+^-chelating agent TPEN (30 µM) or untreated for 30 minutes at 37°C. Afterwards, RNA isolation, cDNA synthesis and qRT-PCR were performed as described above.

### Human nasal fluid binding assay

Adherence of different strains to hNF was assessed using a solid phase-binding assay as previously described (37, 39). Pooled nasal secretion samples from human volunteers was purchased from LeeBio (Maryland Heights, MO, USA, 991-13-S), sonicated on ice to homogenize samples, and protein amount determined using the Coomassie Plus™ protein assay reagent (Thermo Scientific). Briefly, 10 µg total hNF protein (in 100 µl PBS) was immobilized on a 96-well polystyrene plate (Sarstedt REF:82.1581.001) by centrifugation (250 x *g*) for 3 min at room temperature, and incubated overnight at 37°C. Plates were gently washed 3 times with 100 µl DMEM, blocked with 100 µl 0.1% BSA in DMEM for 2 hours, then washed again 3 times with 100 µl DMEM. *Spn* strains to be tested were grown to OD_620_ ∼0.4 in TSY- broth, and diluted into DMEM to 2 × 10^4^ CFU / 100µl, and 100 µl aliquoted into each well. The plate was centrifuged (250 x *g* for 3 min), and incubated at 30°C with 5% CO_2_ for 2 hrs. The wells were then washed 19 times with DMEM, and the adherent bacteria were removed by adding 100 µl of 0.001% Triton X100-PBS and incubating for 15 min at room temperature followed by vigorous mixing. The samples were then serial diluted and plated on TSY-agar plates with antibiotics, and incubated overnight at 37°C with 5% CO_2_. Each strain was tested in triplicate, on three different days.

### ELISA for quantifying capsular polysaccharide

A capture ELISA to determine the CPS amount of pneumococcal isolates was performed as described previously (15). Briefly, Immulon 2HB plates (Thermo Scientific) were coated with serotype 4 specific rabbit antiserum (Staten Institut, Denmark) at a dilution of 1:5,000, and incubated overnight at room-temperature. The next day, strains to be tested were grown to OD_620_ of 0.5, and resuspended in PBS. Samples were sonicated on ice for 30 seconds in total (5 sec on, 15 sec off) using the Branson Sonifier 250 (Duty cycle 40%, output control 2). The antibody coated plate was washed 5 times with wash buffer (10 mM Tris, pH 7.4, 0.02% NaN_3_, 150 mM NaCl, 0.05% Brij), and [100 µl] of two-fold serially diluted bacterial lysate was aliquoted and incubated for 2 hr at RT. The cell lysate was removed, wells washed 3 times with wash buffer, incubated with 1% BSA in PBS for 1 hr, and washed again 3 times. Then [100 µl] [volume] of CPS antibody (1:3000 in 1% BSA/PBS; monoclonal against serotype 4 CPS, kindly provided by Moon Nahm, UAB) was applied and incubated for 24 h with gentle agitation at 4°C. The plates were then washed 5 times with wash buffer, and incubated with goat anti-mouse antibody conjugated to alkaline phosphatase (Sigma-Aldrich; A3688-1 MI; 1:10,000 1% BSA/PBS) for 2 hr at RT. The plate was washed 5 times before being developed with phosphatase substrate PNPP (p-nitrophenyl phosphate) (Thermo Scientific, product number 34047) for 20 min and read at OD_405_ on a Biotek H1 synergy plate reader.

### Statistical Analysis

All statistical analyses were performed using GraphPad Prism (version 9.0) software (GraphPad Software, Inc., San Diego, CA). Unless otherwise specified, statistical differences were calculated using the Mann-Whitney *U* test (comparing two groups) or the Kruskal-Wallis test with Dunn’s post-analysis (comparing multiple groups).

## Acknowledgements

We are grateful for the members of Zafar laboratory (Wake Forest School of Medicine), and Kimberly Walker (UNC Chapel Hill) for comments on the manuscript. This project was supported by startup funds provided by Wake Forest Baptist Medical Center and U.S. Public Health Service grant (R21AI154047) to M.A.Z.

The authors also wish to acknowledge the support of the Wake Forest Baptist Comprehensive Cancer Center Cancer Genomics Shared Resource, supported by the National Cancer Institute’s Cancer Center Support Grant award number P30CA012197.

---

**Supplemental Table S2.**
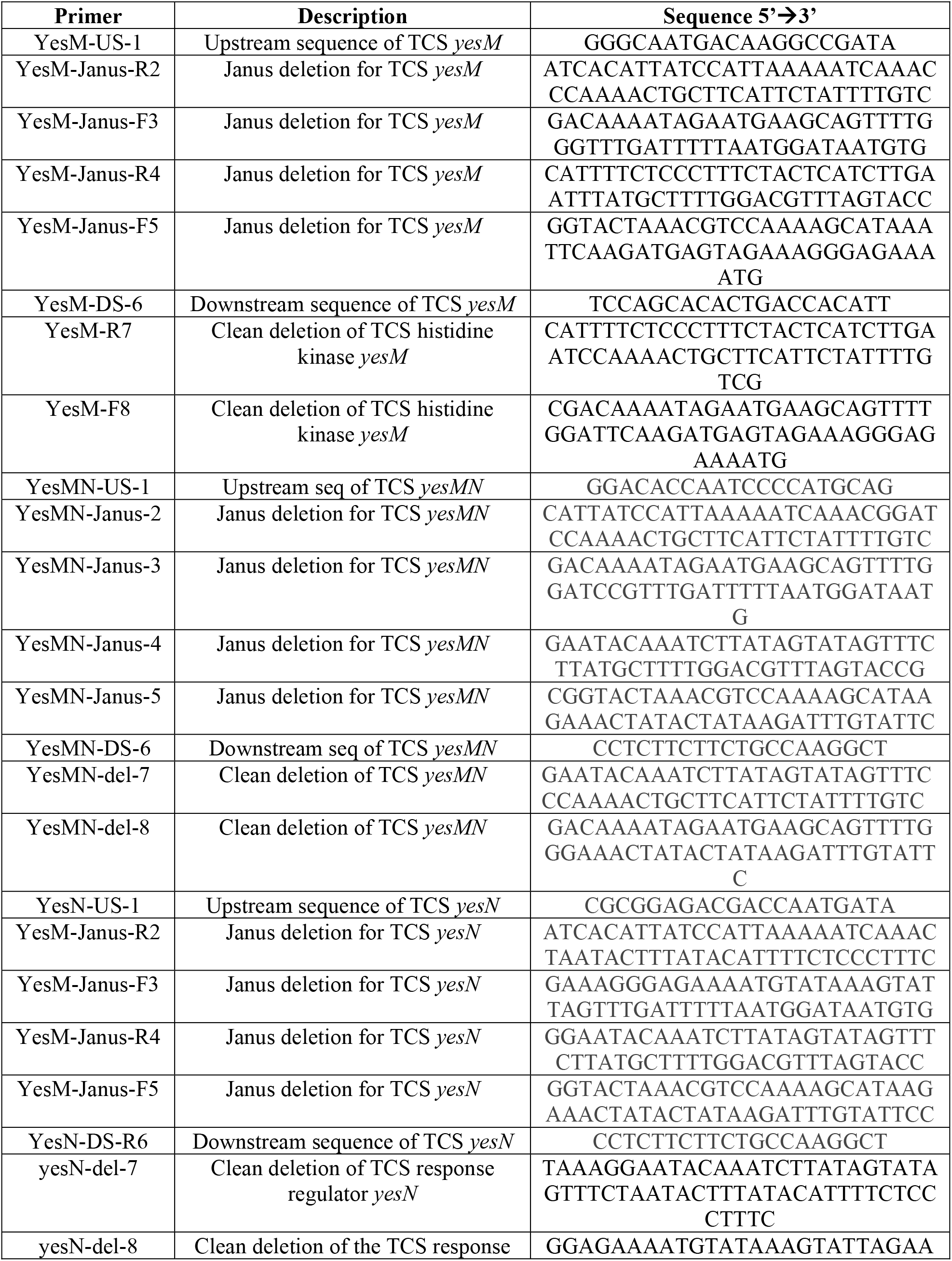
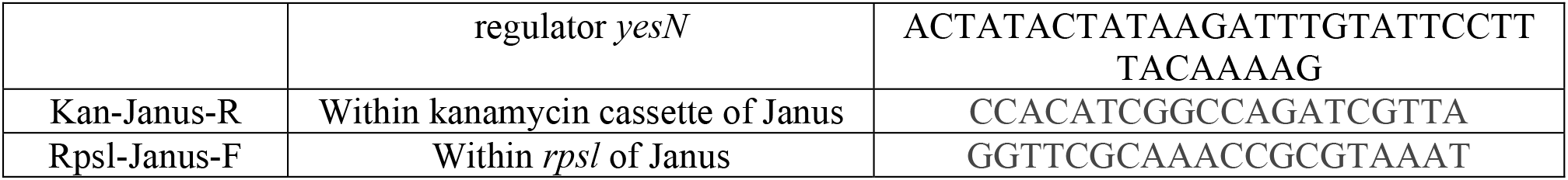
Primers used in the construction of the TCS *yesMN* mutants.

**Supplemental Table S3.**
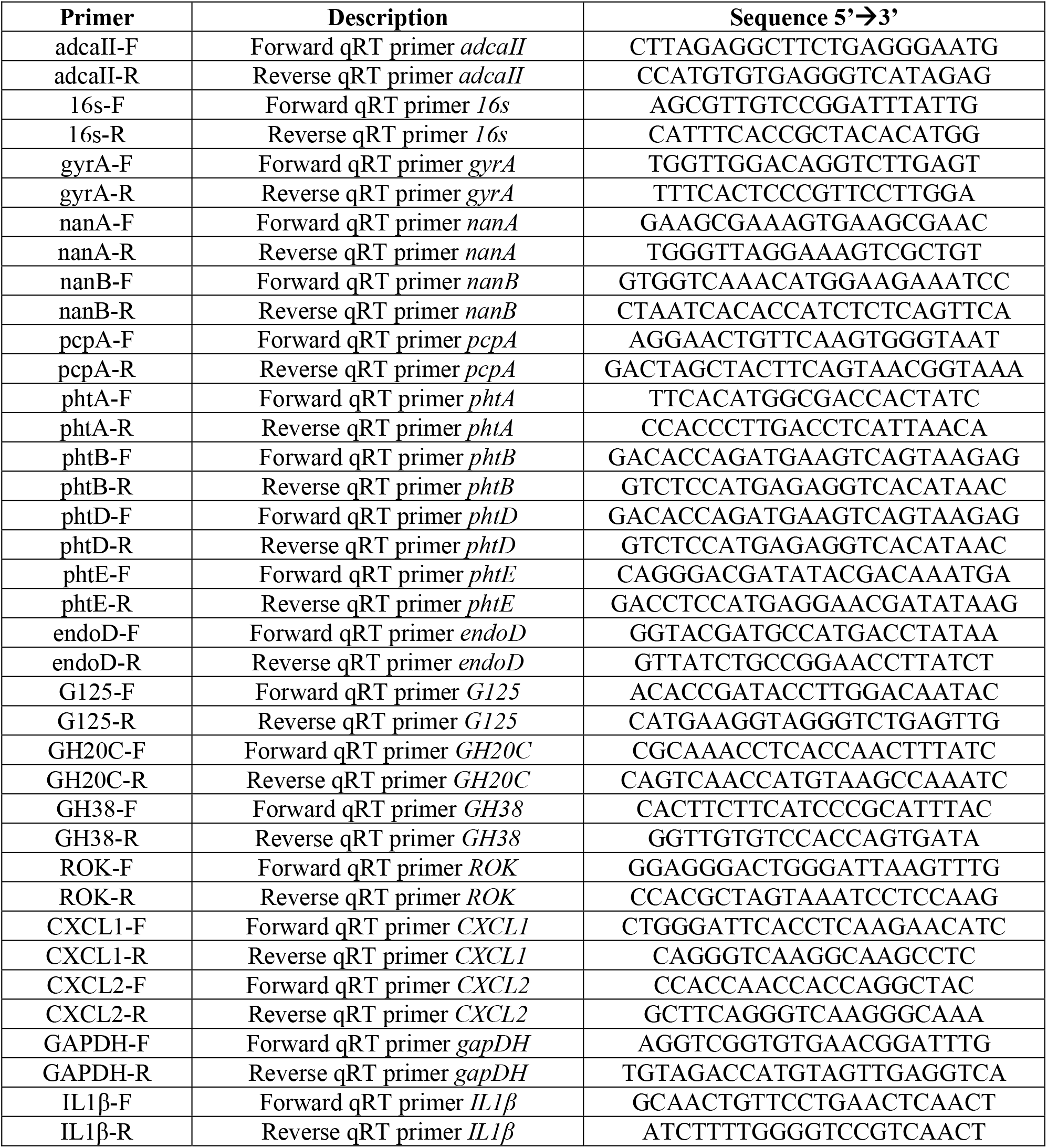
Primers used for qRT-PCR.

**Figure S1.**
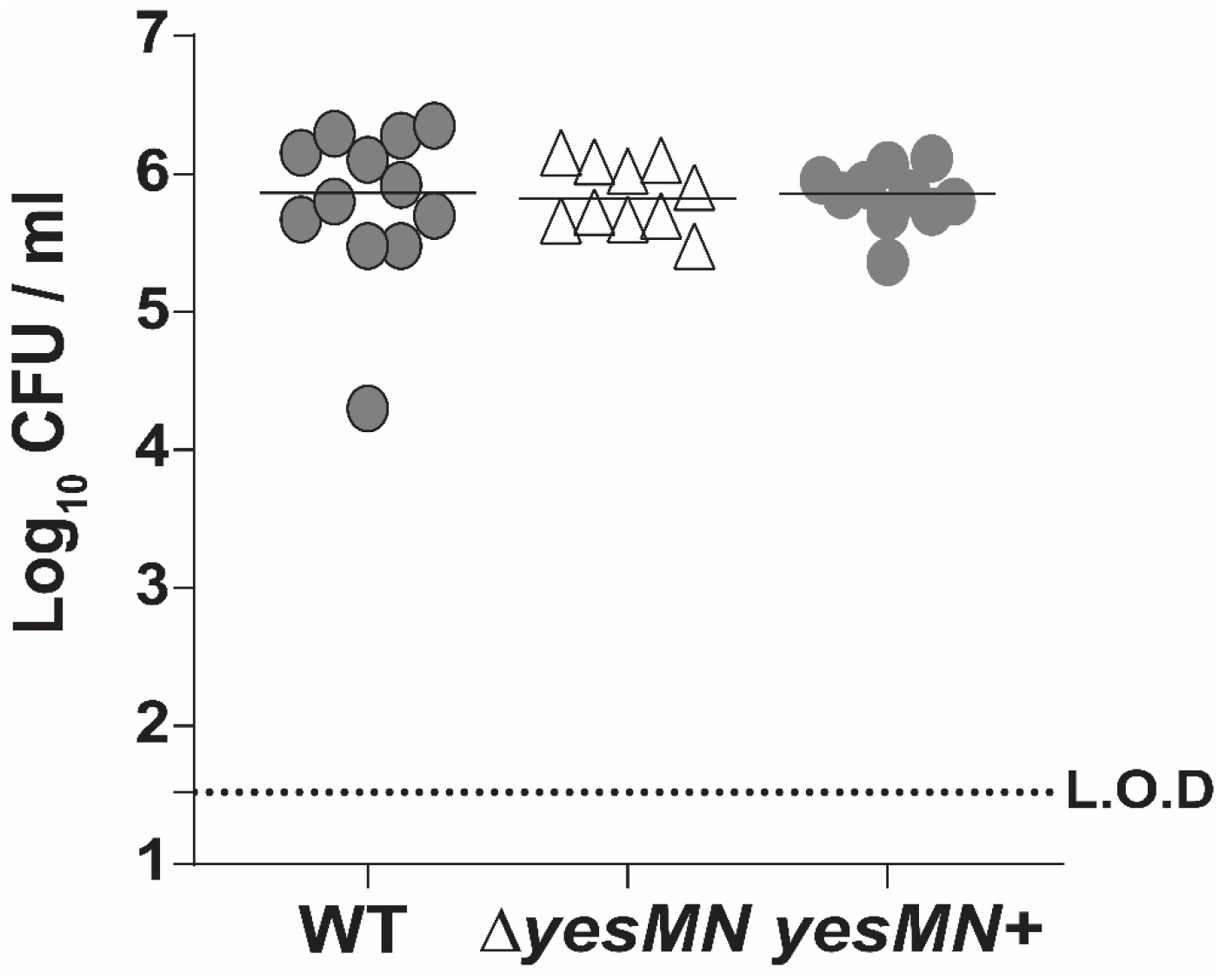
Colonization density of each pneumococcal construct in cultures of URT lavage fluids obtained from pups at day 14 of age. Pups were inoculated i.n at day 4 of life. Median values are shown. Dotted line represents the limit of detection (L.O.D). Statistical differences were calculated using the Kruskal-Wallis analysis of variance with Dunn’s posttest. No statistical differences were observed.

**Figure S2.**
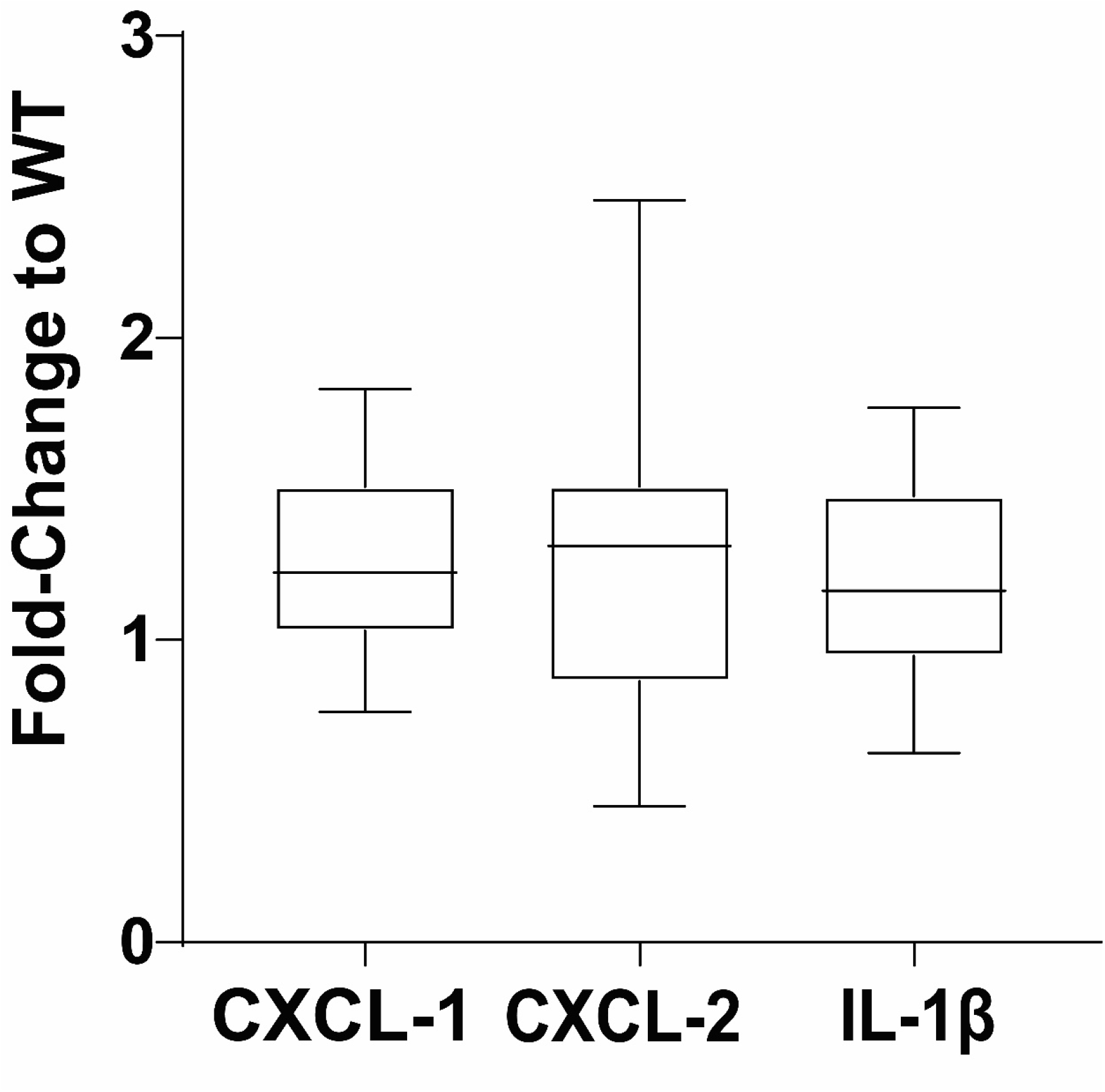
Gene expression as measured by qRT-PCR, for the chemokine/cytokine shown in pups colonized with the *ΔyesMN* mutant relative to that of WT strain inoculated pups at age 7 days. Shown as box and whiskers (Tukey) plot.

**Figure S3.**
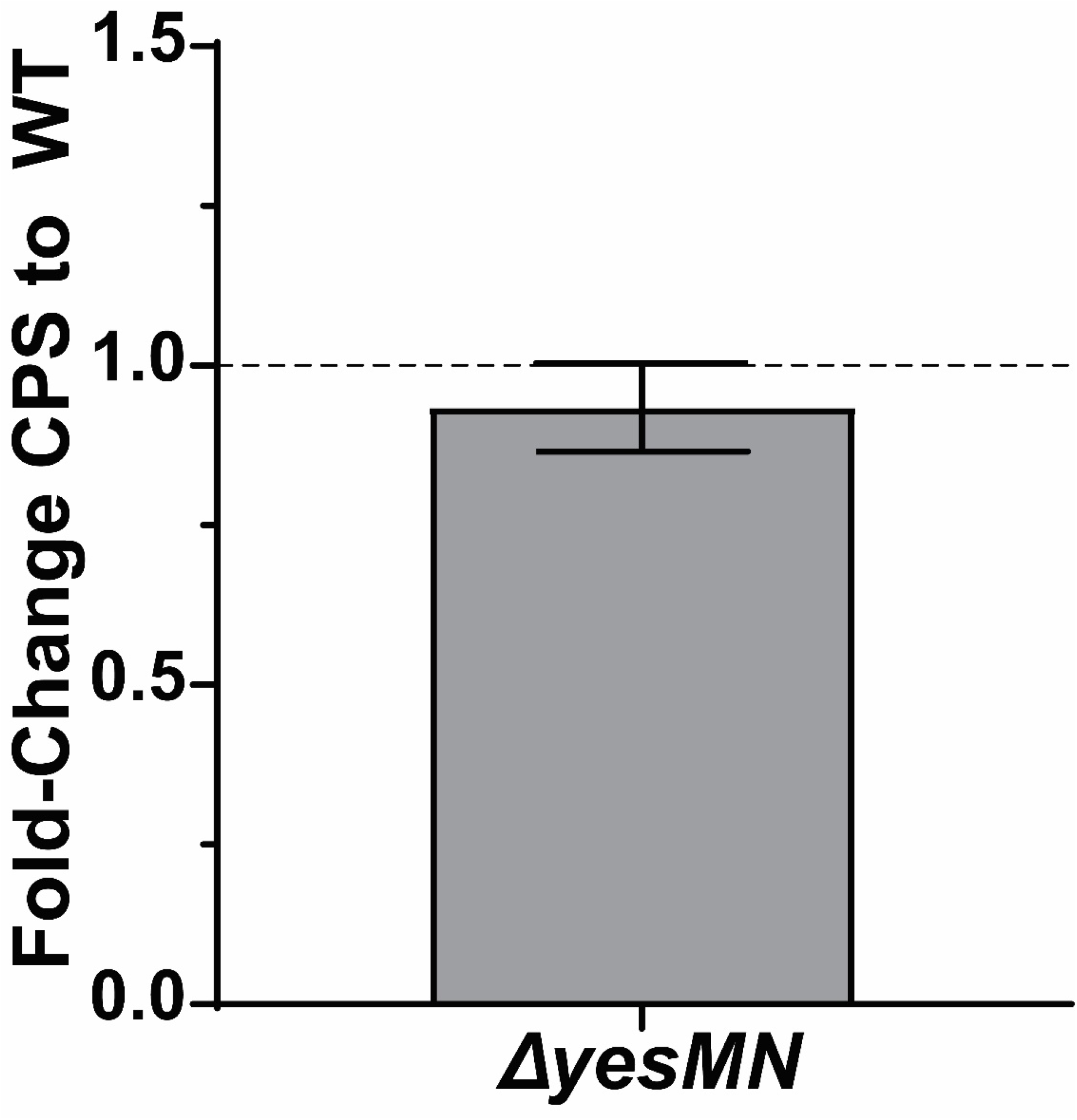
Relative levels of expression of CPS in *ΔyesMN* mutant compared to the WT type 4 isolate using a quantitative capture ELISA. Shown as Mean ± standard errors of mean (SEM).

